# Rapid immunostaining and high-resolution three-dimensional light-sheet microscopy of intact calcified tissues

**DOI:** 10.64898/2026.07.04.736531

**Authors:** Zhangfan Ding, Yixin Shi, Hanyu Liu, Chunjie Li, Junyu Chen, Martine Cohen-Solal, Anjali P. Kusumbe

## Abstract

High-resolution 3D imaging is an important strategy for visualizing and analysing complex skeletal tissue architecture and the bone marrow microenvironment. However, multicolor immunolabeling and imaging of intact skeletal tissues are technologically challenging. The current immunolabeling and clearing methods for intact skeletal elements are very limited, time-consuming and generate low-resolution data or depend on the use of reporter mice. Here, we describe a protocol for efficient clearing and immunolabeling of intact calcified tissues that enables superfast, single-cell resolution, and quantitative 3D light-sheet imaging of intact skeletal elements and teeth. A key aspect of our protocol is the addition of a collagenase digestion step after fixation and decalcification. This step enhances antibody penetration, resulting in deep, comprehensive staining throughout immunostained bones and other calcified tissues. The protocol includes soft tissue removal, fixation, decalcification, bone dehydration, and bleaching, followed by antigen retrieval and permeabilization before the collagenase digestion step. This procedure is performed to prepare the samples for the tissue clearing process that improves bone tissue transparency prior to light-sheet imaging. The entire protocol, from bone collection to image analysis and quantification, takes about 4 days to complete, thus offering significant improvements over previous methods. This protocol is broadly applicable to the visualization of bone microstructure, bone marrow analysis, vascular and neural network mapping, and the study of signaling molecules in bone development and growth. The protocol requires experience with standard tissue processing and immunostaining techniques, and prior experience in tissue clearing and light-sheet imaging is beneficial but not essential.

**Key points:**

- A protocol for efficient clearing and immunolabeling of intact calcified tissues that enables superfast, high-resolution, and quantitative 3D imaging of various intact bones and teeth.
- The entire protocol takes only 4 days to complete the comprehensive staining and perfect transparency throughout the intact bones, offering significant improvements over previous methods.

**Key references:** Biswas, L. et al. *Cell* **186**, 382-397.e24 (2023): https://doi.org/10.1016/j.cell.2022.12.031

## Introduction

Advanced imaging approaches are providing critical insights into the mechanisms involved in highly complex biological processes, such as organ development, regeneration, repair, and pathology^1–3^. These strategies are particularly indispensable for obtaining structural and spatial information and expanding our understanding of crosstalk between vascular, immune, stromal, and neural cells in a tissue context^4–6^. Furthermore, these methods are critical for studying the organization of blood and lymphatic vessels as they enable volumetric rendering across different tissues and organs^7–11^.

Historically, different cell types within tissues have been analyzed in tissue sections using paraffin-embedded sections or thin cryosections. These techniques suffer from high autofluorescence and poor preservation of tissue morphology and epitopes^7,8,12,13^. Additionally, most of these techniques detect only one or two markers and provide very limited 3D information, thus limiting the depth of insights^14^. Capturing 3D information of large tissue areas and volumes at a high resolution is indispensable for understanding the tissue architecture and unraveling multifaceted interactions between different cell types *in situ*^15,16^. Some of the recent studies on thick tissue sections have provided spatial relationships and interactions between different cell types, such as blood vessels, bone cells, and stromal cells^17–20^. However, the sectioning process can lead to distortion in tissue morphology and hinder the accurate reconstruction of large volumes^21–23^, and staining combination is always limited, as recognized by the latest study^20^. It follows that it is far from enough to visualize complex structures such as lymphatic vessels or neurons, which require 3D analysis and may be missed in thin and minute tissues^14,24^.

## Development of the protocol

High-resolution 3D imaging using light-sheet fluorescence microscopy of intact bones has emerged as a promising technique that can offer a deeper understanding of the organization and functions of physiological or pathological systems, organs, and tissues^25–28^. However, applying immunolabeling and 3D imaging to intact skeletal tissues presents significant technical challenges due to their calcified and mineralized nature, limiting the penetration of light and the depth of imaging, especially in thick bone samples^29,30^. Furthermore, preparing bone samples for light-sheet imaging often involves the removal of soft tissues, which can be time-consuming and technically challenging. Failure to remove adjacent tissues adequately can result in suboptimal imaging and a debased visual representation of the anatomical structure of bones. Moreover, achieving uniform and deep immunolabeling within calcified tissue can be problematic and most of the current light-sheet imaging of intact bones relies on the use of reporter mouse lines. Therefore, developing efficient protocols for antibody penetration is crucial for successful imaging.

To address these technical challenges, we recently developed an efficient and robust pipeline for the rapid clearing and immunolabeling of intact calcified tissues that enabled fast, single-cell-level, quantitative 3D light-sheet imaging. Since the fact that the calcified nature impedes the achievement of intact bone tissue clearing and whole-organ immunostaining, we have used this newly-developed pipeline to reveal the lymphatic vessels in bone, which are low-density and extremely difficult to trace their appearance over the past decades, and the role of lymphatic vessels in bone regeneration following injury has also been disclosed with the help of this powerful protocol^31^. This innovative method allows for ultrafast multicolor immunolabeling and clearing of calcified tissues within just 4 days, significantly accelerating the pace of discoveries compared to other imaging pipelines, as shown in the schematic **(Fig. 1).** Here, we describe a detailed protocol for the rapid clearing and immunolabeling of intact bone. The protocol includes a pivotal addition of a collagenase digestion step after fixation and decalcification. This step enhances the penetration of antibodies, resulting in deep and comprehensive staining throughout the immunostained bones and other calcified tissues.

**Fig. 1.**
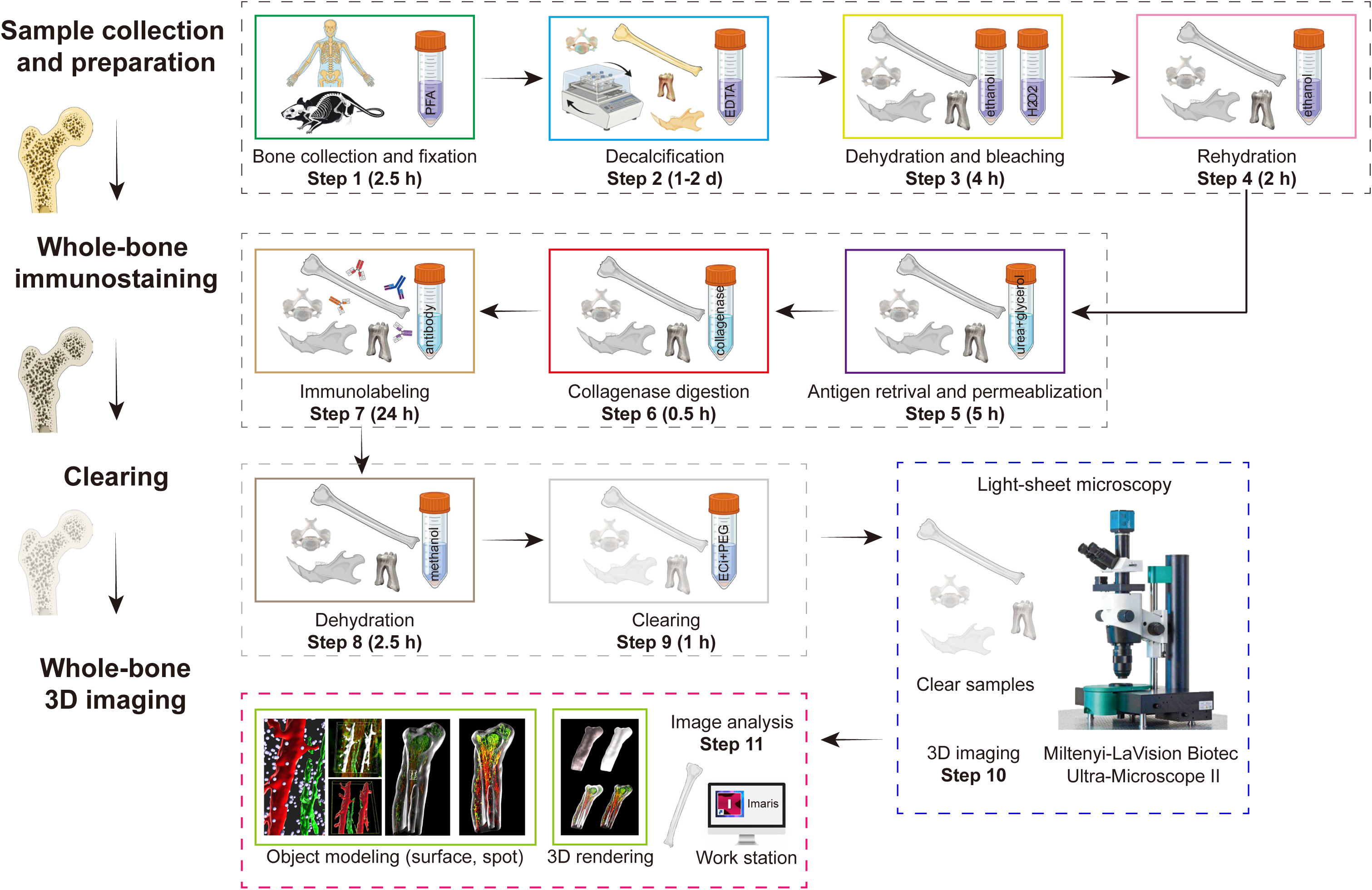
Schematic representation showing the immunostaining and light-sheet imaging method. **Steps 1-4**, Schematic depicting the process of sample collection and preparation. **Steps 5-7**, Timeline of the whole-bone immunostaining for the imaging of light-sheet microscope. **Steps 8-11**, Procedure of tissue clearing strategy, image acquisition and analysis.

In brief, this is a multistep protocol that is easily implementable and scalable. The protocol is initiated with the collection of the whole bone sample, followed by steps that ensure that: (1) surrounding soft tissue is completely removed (Step 1; **Fig. 1**); (2) the bone structure is preserved, while the mineral components and excess water are removed and the bone is bleached (Steps 2-3; **Fig. 1**); (3) bone is prepared for immunolabeling, with a special attention given to the additional step of collagenase digestion after fixation and decalcification, which ensures deep penetration of antibodies (Step 6; **Fig. 1**), thus enabling high-quality imaging of the whole bone; (4) sample is immunostained and thoroughly dehydrated to achieve enough transparency for light-sheet imaging (Steps 7-9; **Fig. 1**); (5) light-sheet microscope is set up properly, and 3D image data are processed and analyzed correctly (Step 10; **Fig. 1**). Collectively, these steps take less than 4 days to complete, and no toxic clearing medium is involved in this procedure. This protocol proposes multiple critical processes to improve the permeability of calcified tissues and to accelerate the realization of tissue clearing, especially the complete removal of the soft tissue around the bone and the adoption of collagenase digestion.

## Applications of the method

Bone immunolabeling combined with light-sheet imaging offers a powerful tool for investigating various aspects of bone biology and pathology. For example, with the optimized protocol, we can obtain samples that allow us to visualize bone microstructure, such as tracing multiple types of cells, including osteocytes, osteoblasts, and osteoclasts and cellular interactions in the bone marrow microenvironment^32^. This enables the study of bone remodeling processes and cell interactions within the bone matrix. A variety of calcified tissues, including mouse tibia, femur, vertebra, hip bone, mandible, maxilla, and tooth, can be cleared through this protocol, and these cleared samples can whereafter be used for bone marrow microenvironment analysis.

This protocol is also compatible with vascular and neural network mapping. Using this approach, multiple murine skeletal elements, including the tibia, vertebra, mandible, and metacarpal bones, can be labeled and visualized at the organ level **(Fig. 2a-f, Supplementary video 1,2)**, providing a valuable framework for studying bone vascularization, angiogenesis, and the role of blood supply in maintaining bone health and facilitating repair. Further, numerous vascular details in whole teeth are revealed by this imaging technique **(Fig. 2g, Supplementary video 3)**. Notably, this approach also enables visualization of the skeletal lymphatic vasculature **(Fig. 2a,b, Extended Data Fig. 1a)**, which is essential for elucidating the role of the lymphatic system in bone homeostasis and disease^33,34^.

**Fig. 2.**
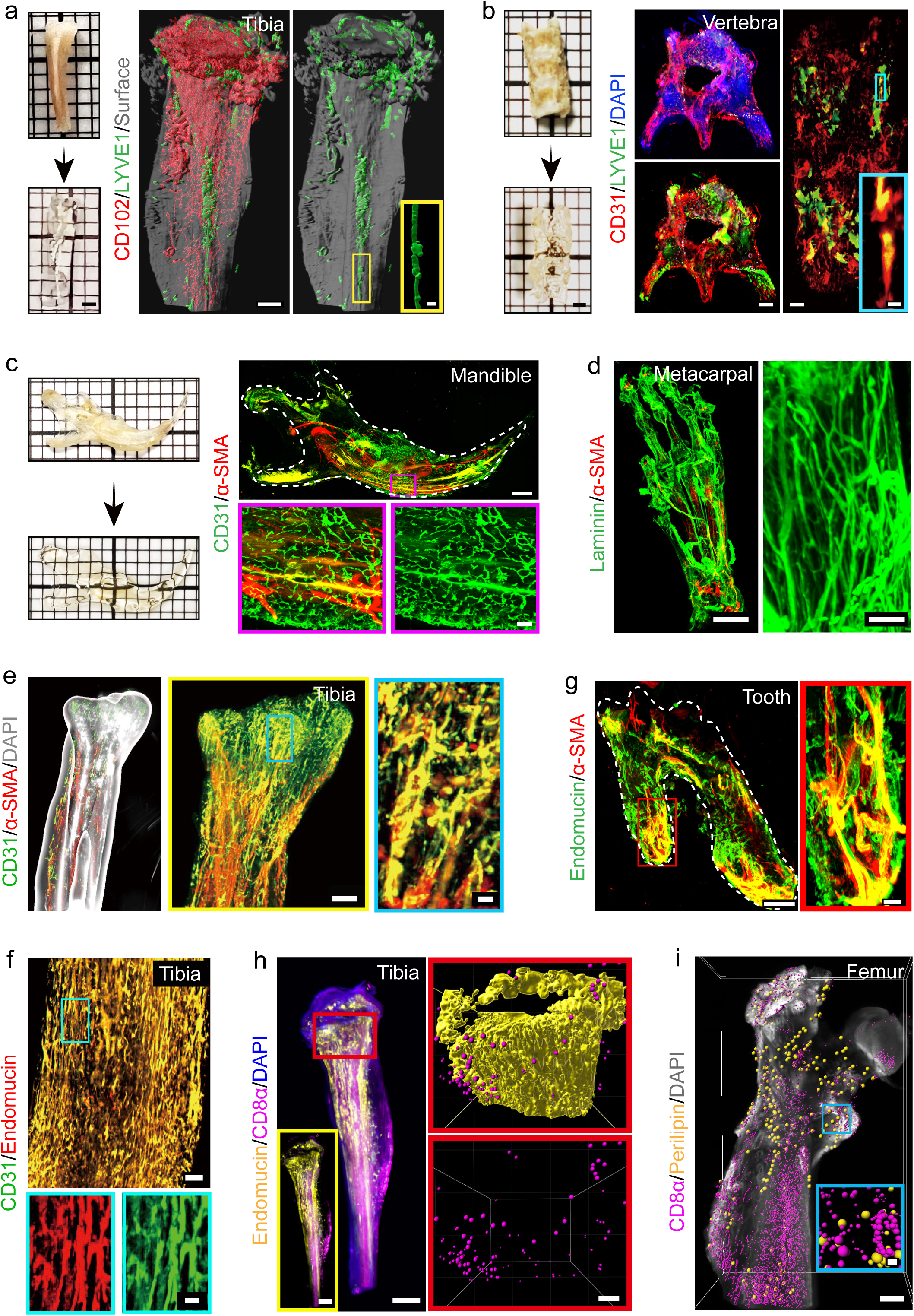
Immunostaining and multicolor panoptic light-sheet imaging for vascular networks and extracellular matrix in bones and teeth. **a**, Images of a mouse tibia prior to and post-tissue clearing. 3D images showing immunostaining for CD102 (red) and LYVE1 (green) with a surface overlay (grey). Inset showing a lymphatic vessel. Scale bars: left, 2 mm; middle, 600 µm; right, 90 µm. **b**, Mouse vertebra prior to and post-tissue clearing. Representative 3D images of a murine vertebra with a high-magnification inset labeled with CD31 (red), LYVE1 (green), and DAPI (blue). Scale bars: left, 2 mm; middle, 500 µm; right, 500 µm; inset, 50 µm. **c**, Mouse mandible prior to and post-tissue clearing. Representative image of a murine mandible showing immunostaining for CD31 (green) and α-SMA (red). High-magnification insets showing the selected region of interest. Scale bars: left, 2 mm; upper, 600 µm; lower, 50 µm. **d**, 3D images of murine metacarpal showing immunostaining for Laminin (green) and α-SMA (red). Inset displaying the selected region of interest. Scale bars: left, 600 µm; right, 90 µm. **e**, Images of diaphysis of mouse tibia obtained from a light-sheet microscopy with high-magnification insets immunostained for CD31 (green) and Endomucin (red). Scale bars: upper, 300 µm; lower, 100 µm. **f**, 3D images of a murine tibia labeled with CD31 (green), α-SMA (red), and DAPI (grey). Insets displaying the blood vessels in the metaphysis. Scale bars: left, 800 µm; middle, 500 µm; right, 200 µm. **g**, Mouse molar prior to and post-tissue clearing. 3D image of a murine molar tooth showing staining for Endomucin (green) and α-SMA (red). Inset showing the selected region of interest. Scale bars: left, 600 µm; right, 90 µm. **h**, 3D images of a mouse tibia acquired on a light-sheet microscopy platform post-clearing with vascular surface reconstruction (yellow) and spot rendering of CD8α-positive T cells (violet). Scale bars: panoptic tibia, 1000 µm; inset, 300 µm. **i**, Representative light-sheet images of a murine femur labeled with CD8α (violet), Perilipin (yellow), and DAPI (grey) generated by 3D reconstruction. Scale bars: panoptic femur, 500 µm; inset, 100 µm.

Additionally, this protocol achieves immunostaining of the whole bone, making it possible to track the distribution of growth factors, signaling molecules, extracellular matrix, and markers of bone development and growth, thus facilitating the study of bone development, growth, and diseases, such as osteoporosis, osteoarthritis, and bone cancers. It enables deep tissue immunolabeling of intact bones, ensuring that the immunolabeled signals remain uniform throughout the entire depth of the sample, as demonstrated by the slice views of the longitudinal layers from different directions **(Extended Data Fig. 1a,b)**. Importantly, this protocol is substantially faster than the current state-of-the-art, producing imaging data in approximately 4 days. Given this speed, the protocol is compatible with drug screening assays to assess the effects of pharmaceutical compounds on bone tissue, as well as investigating the integration of implanted materials.

Lastly, this newly developed protocol is well-suited for comparative anatomical studies and cross-species imaging of bone microstructures, offering insights into evolutionary adaptations and diversity in bone biology. In summary, it enables rapid preparation of intact bone samples for light-sheet imaging with efficient and uniform immunostaining penetration, providing a robust and spatially precise framework for investigating bone biology, health, disease, and regeneration.

## Comparison with other methods

Recent advances in tissue clearing methods, light-sheet microscopy and big data analysis have made it possible to image large tissue volumes at single-cell and nearly subcellular resolutions^35–38^. However, current protocols for immunostaining and whole organ clearing are time-consuming, elaborate, expensive, and organ-specific^39–42^. They also often require the use of either transgenic mice expressing fluorescent reporters and/or toxic and/or corrosive chemicals that might be harmful to the researcher or damage the microscope^43,44^. The requirement for mice expressing fluorescent reporters is particularly true for the calcified tissues, as very few of the existing protocols enable high-resolution multiplex antibody staining of bone tissues. Furthermore, hard tissues make up over 15% of total body weight and are exceptionally challenging to clear^26,45,46^.

Several representative clearing methods have been developed for mineralized tissues over the past decade. Bone CLARITY enables the visualization of intact bone at the cellular level, while its hydrogel-based workflow involves acrylamide, which requires careful handling due to potential neurotoxicity and carcinogenicity^25^. BoneClear supports whole-bone immunolabeling and imaging, whereas the workflow remains relatively lengthy^42^. Osteo-DISCO is useful for visualizing skeletal nerves and vasculature, but it is mainly suited for thin bone samples and reporter-based imaging^47^. PEGASOS and TESOS strategy provide robust solvent-based approaches for calcified tissues and whole-mouse clearing, achieving high transparency and deep volumetric imaging^26,48^. So far, few protocols are available to achieve whole-bone immunostaining and high-resolution imaging in a short time under relatively simple conditions^25,49–52^.

In comparison, our protocol does not depend on the use of reporter mice, which provides flexibility to label multiple targets and select different samples, and comprehensive immunostaining and uniform optical transparency can be achieved within 4 days. Further, the protocol adopts low-cost reagents and an ECi-based clearing system, making it suitable for routine laboratory use. For example, using our protocol, we can immunolabel and image both the bone marrow and intact bone cells. Moreover, the method enables precise immunolabeling of specific target proteins and cell types within bone tissue, such as immune cells and adipocytes, facilitating the visualization of cellular and molecular details within the bone marrow microenvironment **(Fig. 2h,i, Supplementary video 4,5)**. The protocol is compatible with the use of light-sheet microscopy for both rapid and extended imaging sessions, which allows us to capture high-resolution images with high-magnification objectives over longer durations and reveal intricate details of calcified tissue in intact bones, thick bone sections, and whole teeth. In contrast to available protocols which are organ-specific, our protocol applies to a variety of calcified tissues, including long bones, flat bones, irregular bones, and teeth. Furthermore, our approach is not limited to murine samples and can also be applied to human bones, thus broadening its potential applications showed in our recent study^31^. Importantly, one of the most significant advantages of our protocol is a remarkably short processing time for clearing and immunostaining that ensures efficient and timely experimental outcomes under safer working conditions.

## Experimental design

### Method for bone collection and preparation for light-sheet imaging

This protocol provides an efficient, easily implementable and generally applicable method for the collection and preparation of bones for light-sheet imaging. By meticulously removing soft tissue, followed by proper fixation and decalcification, this technique ensures accurate bone imaging. It is specifically designed for cortical bone or bone marrow analysis, making it suitable for various research applications.

### Soft tissue removal (Step 1)

Thoroughly remove soft tissue from freshly dissected bones before fixation. Employ a two-step process involving scraping and collagenase digestion to eliminate adjacent muscle and fat tissue, which may obstruct whole bone imaging. Properly clearing muscle and fat tissue facilitates rapid clearing and yields precise bone structure and shape for imaging. Keep in mind that this technique is ideal for cortical bone or bone marrow analysis but not the periosteum due to possible periosteum damage resulting from scraping and digestion.

### Fixation (Steps 2-3)

Ensure optimal preservation of bone structure by performing proper fixation. Wash the cleaned bones with phosphate-buffered saline (PBS) to eliminate debris and contaminants. Utilize PBS to prepare the fixative solution by combining 4% (wt/vol) paraformaldehyde (PFA) and 0.05% (vol/vol) glutaraldehyde in ice-cold PBS. Prepare the fixative solution immediately before use. Submerge the cleaned bones in the freshly prepared fixative solution and keep them on ice for 2.5 hours. After fixation, wash the bones with PBS three times for 5 minutes each at room temperature (RT, 25 °C) on a rocker platform. Fixed bones can be stored for up to four days at 4 °C before moving to the next step.

### Decalcification (Steps 4-5)

For optimal imaging, remove mineral components from bones through decalcification. Follow these steps: Prepare a 0.5 M ethylenediaminetetraacetic acid (EDTA) solution at pH 7.4. Immerse the fixed bones in the EDTA solution and incubate with continuous rotation for 24 hours at 4 °C. After decalcification, wash the bones with PBS three times for 5 minutes each at RT on a rocker platform.

### Dehydration and bleaching (Steps 6-8)

Dehydrate the bones using an increasing ethanol gradient: Immerse the bone samples in ice-cold dehydration buffer and incubate them at 50%, 80%, and 100% ethanol for 30 minutes each step. Submerge dehydrated bone samples in 100% ethanol for an additional 30 minutes. Keep dehydration buffers on ice during incubation. If necessary, use 5% (vol/vol) hydrogen peroxide (H_2_O_2_) to bleach dehydrated bone samples for 2 hours. Rehydrate the bones by reversing the ethanol gradient and wash them with PBS three times for 20 minutes each.

### Antigen retrieval and permeabilization (Steps 9-10)

Perform antigen retrieval and permeabilization on fixed and bleached bone tissue samples using the ice-cold solution, named light-sheet buffer 1 (LSB1), containing 25% (wt/vol) urea, 15% (vol/vol) glycerol, 7% (vol/vol) Triton X-100, and 10% (vol/vol) DMSO in double-distilled water (ddH_2_O), for 5 hours at 4 °C.

### Enzymatic digestion (Steps 11-12)

Perform enzymatic digestion with 0.2% (wt/vol) collagenase A in PBS, named light-sheet buffer 2 (LSB2), for 30 minutes at 37 °C on a shaker. We found that a concentration of 0.2% with an incubation time of 30 min improves antibody penetration and imaging depth while maintaining the structural integrity of the tissue **(Extended Data Fig. 2a)**. Wash the samples twice for five minutes each on a rocking platform using a washing buffer consisting of 2% (vol/vol) fetal bovine serum (FBS) in PBS.

### Immunolabeling (Steps 13-19)

Incubate the bone tissue samples in a blocking solution containing 10% (vol/vol) donkey serum (DS), 10% (vol/vol) dimethyl sulfoxide (DMSO), and 0.5% (vol/vol) Triton X-100 in PBS for 20 minutes at 37 °C. Incubate the bone samples with primary unconjugated antibodies or Alexa Fluor-conjugated secondary antibodies in a dilution buffer, overnight or 14-16 hours in a water bath at 37 °C, shaking at 70 rpm. Antibody dilution buffer contains 2% (vol/vol) DS, 10% (vol/vol) DMSO, and 0.5% (vol/vol) Triton X-100 in PBS. Wash the bone tissue samples with light-sheet buffer 3 (LSB3), which contains 2% (vol/vol) DS and 0.5% (vol/vol) Triton X-100 in PBS in a 37 °C warm water bath, shaking at 70 rpm, for 3 hours with buffer changes every 15 minutes during the first hour and every 30 minutes for the remaining time. Incubate the samples for 6 to 8 hours at 37 °C in a shaking water bath (70 rpm) with secondary antibodies diluted in the antibody dilution buffer. Incubate the samples with DAPI (1:500) for an additional 1-2 hours. Perform further washing with light-sheet buffer 4 (LSB4) containing 0.5% (vol/vol) Triton X-100 in PBS, in accordance with the washing protocol after primary antibody staining.

### Dehydration (Steps 20-22)

Begin by treating immunostained bone samples with an increasing gradient of ethanol. Immerse the samples successively in 30% ethanol, 50% ethanol, and 80% ethanol solutions at 4 °C, with each step lasting 30 minutes. This gradual ethanol treatment aids in the removal of water from the samples while preserving their structural integrity. After the ethanol treatment, transfer the bone tissues to a container filled with 100% methanol. Incubate the tissues in 100% methanol for a duration of 1 hour at 4 °C with methanol changed every 20 minutes during this duration. This step ensures thorough dehydration of the samples and prepares them for the subsequent clearing process. Following the methanol incubation, rinse the bone tissues with ethyl cinnamate (ECi). Perform two consecutive rinses, each lasting 5 minutes. This rinsing step helps remove any residual ethanol and methanol and prepares the tissues for the clearing solution.

### Tissue clearing (Step 23)

Prepare a clearing solution by combining 80% (vol/vol) ECi and 20% (vol/vol) polyethylene glycol (PEG). This solution serves as the clearing agent, enhancing tissue transparency and enabling optimal imaging. Ensure thorough mixing of the components to create a homogenous solution. Submerge the bone samples in the prepared clearing solution. Allow the bones to incubate in this solution for a period of 30 to 60 minutes. During this incubation, maintain careful rotation of the container to ensure even and complete penetration of the clearing agent. This step is crucial for rendering the bone tissues transparency, facilitating high-quality imaging with light-sheet microscopy.

### Light-sheet imaging setup and image acquisition (Steps 24-29)

Cleared samples were subjected to imaging using the Miltenyi-LaVision Biotech Ultra Microscope II, and image acquisition was facilitated by the LaVision BioTech ImSpector software (MACs, Miltenyi Biotech). The microscope was equipped with a high-quality 2X objective lens designed for the zoom body, featuring a manual zoom range spanning from 0.63X to 6.3X. Importantly, this objective lens was fitted with a Dipping Cap (5.7 mm), which incorporated correction optics optimized for the Olympus MVPLAPO 2X objective. To ensure comprehensive image acquisition, the microscope was outfitted with laser lines at 405, 488, 561, 639, and 785 nm. These laser lines enabled the capture of images across various wavelengths, enhancing versatility in sample analysis. The imaging process was executed using the Neo sCMOS camera manufactured by Andor, known for its superior imaging capabilities. To prepare the samples for imaging, they were meticulously affixed to the sample holder adapter with a drop of superglue. This step ensured stable and secure positioning of the samples during the imaging process. Once securely attached to the sample holder, the samples were gently immersed in ECi contained within a quartz glass cuvette. This immersion process allowed for the optimal exposure of the samples to subsequent light sheet excitation. The imaging process involved exciting the samples with light sheets of varying thickness, ranging from 30 mm to 90 mm, depending on the size of the specific organ of interest. These light sheets were generated at different wavelengths, specifically at 405 nm, 488 nm, 561 nm, 640 nm, and 785 nm, providing a comprehensive spectrum for data acquisition.

### Data conversion and analysis (Steps 30-33)

Raw image data, captured in individual TIFF format, were efficiently converted using the Imaris File converter (version 9.9.0, Bitplane). This conversion process prepared the data for subsequent analysis. The converted image data were then subjected to in-depth analysis using the Imaris software (version 9.9.0, Bitplane). This powerful software enabled precise processing, visualization, and interpretation of the acquired image data, facilitating comprehensive scientific insights.

### Limitations of the method

The availability of suitable light-sheet microscopes and related equipment can be a limiting factor, especially given that setting up and maintaining a light-sheet imaging system can be expensive. Managing and processing the large datasets generated by light-sheet imaging can be computationally intensive. Further, due to the limited scanning area, the light-sheet microscope could only acquire images for whole bones with a small size, such as mouse bone (less than 3 cm in length). For larger cleared samples, subsequent sectioning or trimming into thick slices is often required for individual imaging and three-dimensional reconstruction, which adds workflow and analysis complexity.

## Materials

### Biological materials

C57BL/6 mice (Charles River, Strain#632) between 8-12 weeks were used for general experiments. Six-week-old and 70-week-old mice were used for young and aged studies, respectively.

**▲ CAUTION All animals were maintained following Principles of Laboratory Animal Care formulated by the National Society for Medical Research and the Guide for the Care and Use of Laboratory Animals (National Academies Press, 2011)**

### Reagents

- Phosphate-buffered saline (10×) (PBS; VWR, cat. no. 437117K)
- Paraformaldehyde (PFA; Sigma-Aldrich, cat. no. P6148)

**▲ CAUTION PFA is a highly toxic material; avoid direct contact with skin, eyes or mucous membranes; avoid breathing the powder during measuring and preparation; handle it in a fume hood to avoid breathing it in and contaminating the air**

- Glutaraldehyde (Sigma-Aldrich, cat. no. 340855)

**▴ CAUTION Glutaraldehyde is a toxic chemical; avoid direct contact with skin, eyes or mucous membranes**

- Ethylenediaminetetraacetic acid (EDTA; Sigma-Aldrich, cat. no. E6758)
- Ethanol (Sigma-Aldrich, cat. no.1070172511)
- Hydrogen peroxide (H_2_O_2_; Sigma-Aldrich, cat. no. H1009)
- Triton X-100 (Sigma-Aldrich, cat. no. T8787)

**▲ CAUTION Triton X-100 is hazardous and may cause severe damage to eyes**

- Urea (VWR, cat. no. 28876.367)
- Glycerol (VWR, cat. no. 24388.260)
- Collagenase A (Merck, cat. no. 10103578001)
- Fetal bovine serum (FBS; Sigma-Aldrich, cat. no. F7524)
- Donkey serum (DS; Abcam, cat. no. ab7475)
- Dimethyl sulfoxide (DMSO; Sigma-Aldrich, cat. no. D5879)

**▲ CAUTION DMSO is hazardous and may penetrate the skin and cause damage**

- Methanol (ThermoFisher Scientific, cat. no. M/4058/17)

**▲ CAUTION Methanol is hazardous and may cause severe damage to skin, mucous membranes, respiratory tract, and eyes; handle it in a fume hood to avoid breathing it in and contaminating the air**

- Ethyl cinnamate (ECi; Sigma-Aldrich, cat. no. 112372)
- Polyethylene glycol (PEG; Sigma-Aldrich, cat. no. 447943)
- 4’,6-diamidino-2-phenylindole (DAPI; Sigma-Aldrich, cat. no. D9542)

**▲ CAUTION DAPI is carcinogenic and teratogenic; avoid direct contact with skin, eyes or mucous membranes**

- Superglue (No-Nonsense, cat. no. 26780)

### Equipment

#### Consumables

- 15 mL conical tube (Corning, cat. no. 352096)
- 50 mL conical tube (Corning, cat. no. 352070)
- Pipettes (P10, P20, P200, P1000) (Gilson Pipetman, cat. no. F144802, F123600, F123601, and F123602)
- Pipette tips (P10, P20, P200, P1000) (Eppendorf, cat. no. 0030000811, 0030000854, 0030000870, and 0030000927)
- 5 mL serological pipettes (Corning, cat. no. 4478)
- 10 mL serological pipettes (Falcon, cat. no. 357551)
- Serological pipette gun (ThermoFisher Scientific, cat. no. 10402822)
- Fine-tipped tweezers (Merck, cat. no. T4287-1EA)

#### Common equipment

- Rocking platform (VWR, cat. no. 444-016)
- Water bath (Stuart, cat. no. SBS40)

#### Specific equipment

- Miltenyi-LaVision Biotec Ultra-Microscope II (Miltenyi Biotec)
- LaVision BioTech ImSpector MACs (Miltenyi Biotec)

#### Software

- Imaris (version 9.9.0) (Bitplane, https://imaris.oxinst.com/packages)
- Imaris File converter (version 9.9.0) (Bitplane, https://imaris.oxinst.com/packages)
- Adobe Photoshop CC 2023 (Adobe, https://www.adobe.com/uk/products/photoshop)
- Adobe Illustrator CC 2023 (Adobe, https://www.adobe.com/uk/products/illustrator)
- GraphPad Prism (version 9) (GraphPad, https://www.graphpad.com)

### Reagent setup

#### PBS solution (1×)

Prepare PBS solution (1×) by adding 10 mL PBS solution (10×) to 90 mL double distilled water (ddH_2_O) and autoclave it. PBS can be stored at RT for 1 year.

#### Fixation solution (4% (wt/vol) PFA and 0.05% (vol/vol) glutaraldehyde)

To make 100 mL of fixative solution, add 4 g PFA and 50 µL of glutaraldehyde in 99.95 mL of PBS. Divide the solution into aliquots and store them at 20 °C for up to 4 weeks.

#### Decalcification solution (0.5 M EDTA, pH 7.4)

Dissolve 14.6 g of EDTA in 95 mL of ddH_2_O by adding ∼2 g of NaOH pellets and stirring continuously until the pH approaches 8.0. After the solution has cooled to RT, gradually add 0.1 M HCl together with ddH_2_O while continuously monitoring the pH and total volume, until the pH reaches 7.4 and the total volume reaches 100 mL. In this step, operate and check slowly and repeatedly to ensure accurate concentration and pH value of the decalcification solution. Store it at 4 °C for 6 months.

#### Bleaching solution (1.5% (vol/vol) H_2_O_2_ and 95% (vol/vol) ethanol)

To prepare 100 mL of bleaching solution, add 5 mL 30% H_2_O_2_ and 5 mL DMSO to 90 mL of ethanol. Prepare fresh each time.

#### Light-sheet buffer 1 (LSB1) (25% (wt/vol) urea, 15% (vol/vol) glycerol, 7% (vol/vol) Triton X-100, and 10% (vol/vol) DMSO)

To make 100 mL of LSB1, add 25 g urea, 15 mL glycerol, 7 mL Triton X-100, and 10 mL DMSO to 60 mL ddH_2_O. Place the container on the shaker until the urea has been completely dissolved. After the solution has cooled down to RT, adjust the final volume to 100 mL with ddH_2_O. Keep the buffer at 4 °C. Make up fresh each time and dispose of any unused solution properly.

#### Light-sheet buffer 2 (LSB2) (0.2% (wt/vol) collagenase A)

To make 100 mL of LSB2, dissolve 0.2 g collagenase A in 100 mL PBS. It should be prepared freshly each time.

#### Blocking solution (10% (vol/vol) DS, 10% (vol/vol) DMSO, and 0.5% (vol/vol) Triton X-100)

To prepare blocking solution by adding 10 mL DS, 10 mL DMSO, and 0.5 mL Triton X-100 to 79.5 mL PBS. Prepare fresh each time.

#### Antibody dilution buffer (2% (vol/vol) DS, 10% (vol/vol) DMSO, and 0.5% (vol/vol) Triton X-100)

To make 100 mL of antibody solution, add 2 mL DS, 10 mL DMSO, and 0.5 mL Triton X-100 to 87.5 mL PBS. The dilution ratios of antibodies are listed in **Extended Data Table 1**. The solution should be prepared freshly each time.

#### Light-sheet buffer 3 (LSB3) (2% (vol/vol) DS and 0.5% (vol/vol) Triton X-100)

To make 100 mL of LSB3 by adding 2 mL DS and 0.5 mL Triton X-100 to 97.5 mL PBS and mixing well. Store it at 4 °C for up to 1 week.

#### Light-sheet buffer 4 (LSB4) (0.5% (vol/vol) Triton X-100)

To make 100 mL of LSB4 by adding 0.5 mL Triton X-100 to 99.5 mL PBS and mixing well. Store it at 4 °C for up to 1 week.

#### Clearing medium (80% (vol/vol) ECi and 20% (vol/vol) PEG)

Prepare 100 mL of clearing solution by mixing 80 mL ECi and 20 mL PEG. The solution should be prepared freshly each time.

## Procedure

### Collection of samples

● **TIMING variable**
1 Cull 8-12 week-old mice (C57BL/6J mice) by cervical dislocation to isolate the bones. Rinse the animals with 70% ethanol and make an incision at the peritoneal cavity. Remove the skin by pulling from both sides. Collect all the bones carefully and wash them with PBS.

▲ **CRITICAL STEP** Carefully dispose of the surrounding muscles and fats before fixing to ensure successful whole bone imaging. Employ a two-step process involving trimming off muscles with scissors and eliminating residual soft tissues under a microscope **(Extended Data Fig. 2b)**.

? TROUBLESHOOTING

### Bone fixation

● **TIMING 2.5 h**
2. Wash the freshly collected bones thrice with PBS to remove debris and contaminants. Immerse the cleaned bones immediately into the ice-cold fixation solution of 4% (wt/vol) PFA and 0.05% (vol/vol) glutaraldehyde for 2.5 hours in a 50 mL Falcon tube.

▲ **CRITICAL STEP** Perfusion fixation is not required for this protocol, and the method is also compatible with perfusion-fixed samples. Prepare the fixation solution freshly and keep it ice cold during its use. Maintain the bones in the fixative solution and keep shaking while fixing them at 4 °C. Handle PFA with extreme caution, as it is hazardous, highly flammable, and toxic. The current protocol provides robust detection for the targets examined in this study. However, for certain low-abundance proteins, longer fixation times may improve protein retention, signal preservation, and staining quality. Therefore, fixation duration should be optimized depending on the abundance and properties of the target protein, as well as the experimental application. For imaging of low-abundance proteins, we recommend extending the fixation to the standard duration of 12 hours.

? TROUBLESHOOTING

3. Wash all the samples 3 times with 10 mL PBS at RT for 5 minutes each time. Ensure thorough washing to remove fixative agents.

▲ **CRITICAL STEP** Utilize a rocker platform for gentle agitation during the washing steps. By meticulously following these precise steps, you can ensure proper fixation of bone samples, promoting the optimal preservation of bone structure for your scientific investigations.

▪ **PAUSE POINT** Fixed samples can be stored in PBS for up to 4 days at 4 °C.

### Decalcification

● **TIMING 24-48 h**
4. Submerge bone tissue in a 50 mL Falcon tube containing more than 45 mL of 0.5 M EDTA solution (pH 7.4) and incubate at 4 °C under constant rotation. Incubate bones from young mice for 24 hours and from aged mice for 48 hours.

▲ **CRITICAL STEP** The pH value of the decalcification solution must be 7.4 to obtain the best outcome without tissue damage. Constant rotation is required for homogenous decalcification.

? TROUBLESHOOTING

5. Wash all the samples three times with 5 mL PBS on a rocker. Each wash should last for 5 minutes at 4 °C.

? TROUBLESHOOTING

### Dehydration

● **TIMING 2 h**
6. Prepare the ethanol solution with PBS. Immerse all the bones successively in an increasing gradient of ethanol i.e. 50%, 80%, 100% for dehydration. Incubate the samples in more than 45 mL of each solution in a 50 mL Falcon tube at 4 °C for 30 minutes per concentration.

▴ **CRITICAL STEP** Incubate the sample under gentle rotation. Change 100% ethanol twice after 20 minutes for extended dehydration. Keep the dehydration solution on ice to ensure that it is ice-cold throughout the incubation to minimize its influence on the tissue.

? TROUBLESHOOTING

### Bleaching

● **TIMING 2 h**
7. Prepare the bleaching solution in ethanol. Bleach all the dehydrated bone samples by immersing them in a solution of 1.5% H_2_O_2_ (vol/vol) in ethanol for 2 h in a 50 mL Falcon tube at RT under gentle rotation. This step helps in achieving effective sample bleaching.

▲ **CRITICAL STEP** Avoid aqueous bleaching solutions, as they may cause rapid rehydration and tissue swelling. Ethanol is more compatible with dehydrated tissues and improves reagent penetration and bleaching efficiency.

? TROUBLESHOOTING

### Rehydration

● **TIMING 2 h**
8. Reverse the ethanol gradient to rehydrate the bones. Immerse all the samples with a decreasing gradient of ethanol concentrations at 4 °C in a similar way as in step 6. Wash all the samples thrice with 45 mL PBS for 20 minutes each time after rehydration.

▲ **CRITICAL STEP** Wash the sample under gentle rotation to thoroughly remove residual ethanol and any bleaching agents if used.

### Antigen retrieval and permeabilization

● **TIMING 5 h**
9. Use ddH_2_O to prepare the ice-cold LSB1 containing 25% urea (wt/vol), 15% glycerol (vol/vol), 7% Triton X-100 (vol/vol), and 10% DMSO (vol/vol). Mix them well and store the buffer at 4 °C until use.
10. Submerge the bone samples in a 50 mL Falcon tube containing more than 45 mL of LSB1 solution, ensuring that the tissues are fully submerged, and incubate at 4 °C for 5 hours. This step will facilitate the optimal permeabilization of bone tissue and allow effective antibody penetration.

▲ **CRITICAL STEP** Keep the samples on a rotator to ensure thorough permeabilization.

? TROUBLESHOOTING

▪ **PAUSE POINT** Samples can be stored in PBS for up to 4 days at 4 °C.

### Enzymatic digestion

● **TIMING 0.5 h**
11. Process all the samples in a 50 mL Falcon tube containing more than 45 mL of freshly prepared LSB2 solution consisting of 0.2% collagenase A (wt/vol) in PBS. Ensure that the tissues are fully submerged and incubate at 37 °C for 30 minutes with constant shaking.

▲ **CRITICAL STEP** Prepare LSB2 freshly each time. Place the samples in a shaker to ensure even and efficient digestion.

? TROUBLESHOOTING

12. Wash samples twice for 5 minutes each in 45 mL solution containing 2% FBS (vol/vol) in PBS at 37 °C on a rocker to stop digestion.

### Sample blocking

● **TIMING 20 min**
13. Prepare a blocking solution composed of 10% DS (vol/vol), 10% DMSO (vol/vol), and 0.5% Triton X-100 (vol/vol) in PBS.
14. Incubate the bone samples in a 50 mL Falcon tube containing more than 45 mL of blocking solution for 20 minutes at 37 °C on a shaker. This step helps to prevent non-specific binding of antibodies and enhances signal specificity.

? TROUBLESHOOTING

### Primary antibody incubation

● **TIMING 14-16 h**
15. Prepare the primary antibody dilution buffer composed of 2% DS (vol/vol), 10% DMSO (vol/vol), and 0.5% Triton X-100 (vol/vol) in PBS.
16. Incubate all samples with antibodies (conjugated or primary unconjugated as needed) in a 1.5 mL Eppendorf tube containing approximately 750 μL of antibody solution, ensuring that the tissues are fully submerged. Incubate overnight or for 14-16 hours at 37 °C in a water bath with constant shaking at 70 rpm. The recommended incubation time of different calcified tissues is provided in **Extended Data Table 1**. The sample must be immersed in the antibody solution all the time. All the primary antibodies are listed in **Extended Data Table 1** with their dilution factors for optimal outcomes.

▲ **CRITICAL STEP** Prepare antibody dilution buffer freshly each time. Samples being stained with conjugated primary antibody should be kept from light from this step onward. As described in our previous study^14^, a panel of antibodies validated for fluorescence staining in bone tissues is provided as a useful reference. In general, antibodies that perform well on cryosections are more likely to be compatible with tissue-clearing-based staining. We recommend first screening candidate antibodies on bone tissue sections, followed by validation in whole-tissue clearing and imaging.

? TROUBLESHOOTING

### Washing after antibody incubation

● **TIMING 3 h**
17. Prepare the washing solution, which we named LSB3, composed of 2% DS (vol/vol) and 0.5% Triton X-100 (vol/vol) in PBS. Perform a thorough washing of the bone samples with 10 mL LSB3 in a 37 °C warm water bath for 3 hours, ensuring gentle shaking at 70 rpm.

▲ **CRITICAL STEP** Change LSB3 every 15 minutes for the first hour and then every 30 minutes for the remaining two hours.

? TROUBLESHOOTING

### Secondary antibody incubation

● **TIMING 7-10 h**
18. Incubate samples with fluorescent dye-conjugated secondary antibodies for 6-8 hours in a 1.5 mL Eppendorf tube containing approximately 750 μL of antibody solution, ensuring that the tissues are fully submerged. Incubate at 37 °C in a water bath with gentle shaking at 70 rpm if staining with unconjugated primary antibodies. The recommended incubation time of different calcified tissues is provided in **Extended Data Table 1**. Incubate the samples with DAPI (1:500) for an additional 1-2 hours. All the secondary antibodies are listed in **Extended Data Table 1** with their dilution factors.

▲ **CRITICAL STEP** Prepare the secondary antibody solution as the primary antibody, and prepare it freshly each time. Protect the incubated samples from direct light from this step on. Ready-to-use DAPI solution for cell-based staining are not suitable for staining optically cleared tissues in this protocol. Instead, highly purified DAPI should be used to prepare DAPI solution (1:500) to ensure rapid permeation into deep tissue. Further, longer-wavelength nuclear dyes, such as TO-PRO-3, can also be used as an alternative when deeper penetration or reduced signal attenuation is required.

? TROUBLESHOOTING

### Final washing step

● **TIMING 3 h**
19. Conclude the immunolabeling process by performing further washing steps following the established washing protocol. The washing solution is 0.5% Triton X-100 (vol/vol) in PBS which is regarded as LSB4. Wash samples in a similar way as in step 17 to completely remove the excess antibodies and DAPI. This step ensures the removal of unbound antibodies and maintains the specificity of the labelling.

▲ **CRITICAL STEP** Protect the incubated samples from direct light.

? TROUBLESHOOTING

### Dehydration

● **TIMING 2.5 h**
20. Following immunolabeling, dehydrate all the samples by immersing them in an increasing gradient of ethanol (30%, 50%, and 80%) for 30 minutes in a 50 mL Falcon tube while gently rotating at 4 °C.

? TROUBLESHOOTING

21. Subsequently, immerse tissue samples in 100% methanol for 1 hour at 4 °C under gentle rotation, with the methanol solution changed every 20 minutes during this incubation.

? TROUBLESHOOTING

### Tissue clearing

● **TIMING 1 h**
22. Remove the methanol completely and rinse the samples twice with 5 mL ECi each.
23. Clear the tissue samples by immersing them in the clearing solution composed of 80% ECi (vol/vol) and 20% PEG (vol/vol) in a 50 mL Falcon tube, ensuring that the tissues are fully submerged. Incubate at RT for 30-60 minutes under gentle rotation.

? TROUBLESHOOTING

### Light-sheet imaging

● **TIMING variable Imaging setup**
24. Utilize the Miltenyi-LaVision Biotech Ultra microscope II along with the LaVision BioTech ImSpector software to capture images of cleared tissue samples.

### System initialization

25. Power on the light-sheet microscope and allow the lasers and lamps to warm up. Launch the software and meticulously verify that all settings are configured correctly.

### Objective lens

26. Ensure that the microscope is equipped with a 2X objective lens for zoom functionality, featuring a manual zoom range of 0.63X-6.3X. The objective should be fitted with a Dipping cap (5.7 mm) containing correction optics designed for the Olympus MVPLAPO 2X.

### Laser lines and camera

27. Verify that the microscope includes laser lines at 405 nm-100 mW, 488 nm-85 mW, 561 nm-100 mW, 639 nm-70 mW, and 785 nm-75 mW, as well as a Neo sCMOS camera (Andor).

### Sample attachment

28. Attach the cleared samples to the sample holder adaptor using a small drop of superglue.

▲ **CRITICAL STEP** Ensure that the samples are dry before attaching them to the adaptor. Attach the samples carefully using forceps to prevent damage to the organ. Avoid introducing bubbles during the fixation of samples on the adaptor. The superglue will become effective after 30-60 seconds. Handle sample immersion with care for optimal exposure.

? TROUBLESHOOTING

### Sample excitation

29. Gently immerse the sample in ECi within a quartz glass cuvette. Excite the sample with light sheets at the required wavelengths.

? TROUBLESHOOTING

### Data processing

30. Convert the acquired raw data files using the Imaris File converter (version 9.9.0) and perform subsequent analysis using Imaris software (version 9.9.0).

### Image analysis and quantifications

● **TIMING variable Software utilization**
31. Adopt a workstation to analyze the high amount of data set of acquired images **(Extended Data Table 2)**. Utilize Imaris software (version 9.9.0) to reconstruct and process the Z-sections of the light-sheet images. Apply the animation function of the software to make videos showcasing the spatial visual effect **(Supplementary video 6)**.

### Vascular density quantification

32. Carry out basic 3D analysis of images **(Supplementary video 7)**. Employ surface analysis XTensions tools within Imaris to perform vascular density quantification. Utilize the 3D crop tool to analyze the total tissue volume.

### 3D surface rendering

33. Utilize the 3D surface rendering tool in Imaris to visualize the components of the organ effectively. Define the Regions of Interest (ROI) for individual channels of cells, matrix components, or vessels, and then reconstruct the surface. Smoothen the ROI post-surface reconstruction and subtract the background using appropriate settings and preview the final image **(Supplementary video 8)**.

### Timing

Once all samples are prepared properly, reagents are obtained and set up according to the above requirements and ideal experiment conditions have been identified, the entire protocol, from bone collection (Step 1) to image analysis and quantifications (Steps 31-33), takes ∼4 days to complete. The hands-on timing for each stage of the PROCEDURE is summarized below.

Step 1-3, bone collection and fixation: 2.5 h

Step 4-5, decalcification: 24-48 h

Step 6-7, dehydration and bleaching: 4 h

Step 8, rehydration: 2 h

Step 9-10, antigen retrieval and permeabilization: 5 h

Step 11-12, enzymatic digestion: 0.5 h

Step 13-19, immunolabeling: 24 h

Step 20-21, dehydration: 2.5 h

Step 22-23, tissue clearing: 1 h

Step 24-30, variable

Step 31-33, variable

### Troubleshooting

Troubleshooting advice can be found in **Table 1**.

### Anticipated results

This study presents a rapid multicolor immunolabeling and clearing strategy for intact calcified tissues, including bone and tooth, enabling three-dimensional light-sheet microscopy imaging of whole bones within 4 days **(Fig. 1)**. Although whole-mouse imaging and intravital tissue clearing have recently emerged and attracted considerable attention^48,53,54^, immunostaining and imaging of intact calcified tissues remain difficult to achieve using previously reported protocols owing to either limitations of the equipment or excessive time cost. Using the protocol described here, robust whole-tissue transparency can be achieved across multiple bone samples **(Fig. 2a-c**, **Fig. 3a)**.

**Fig. 3.**
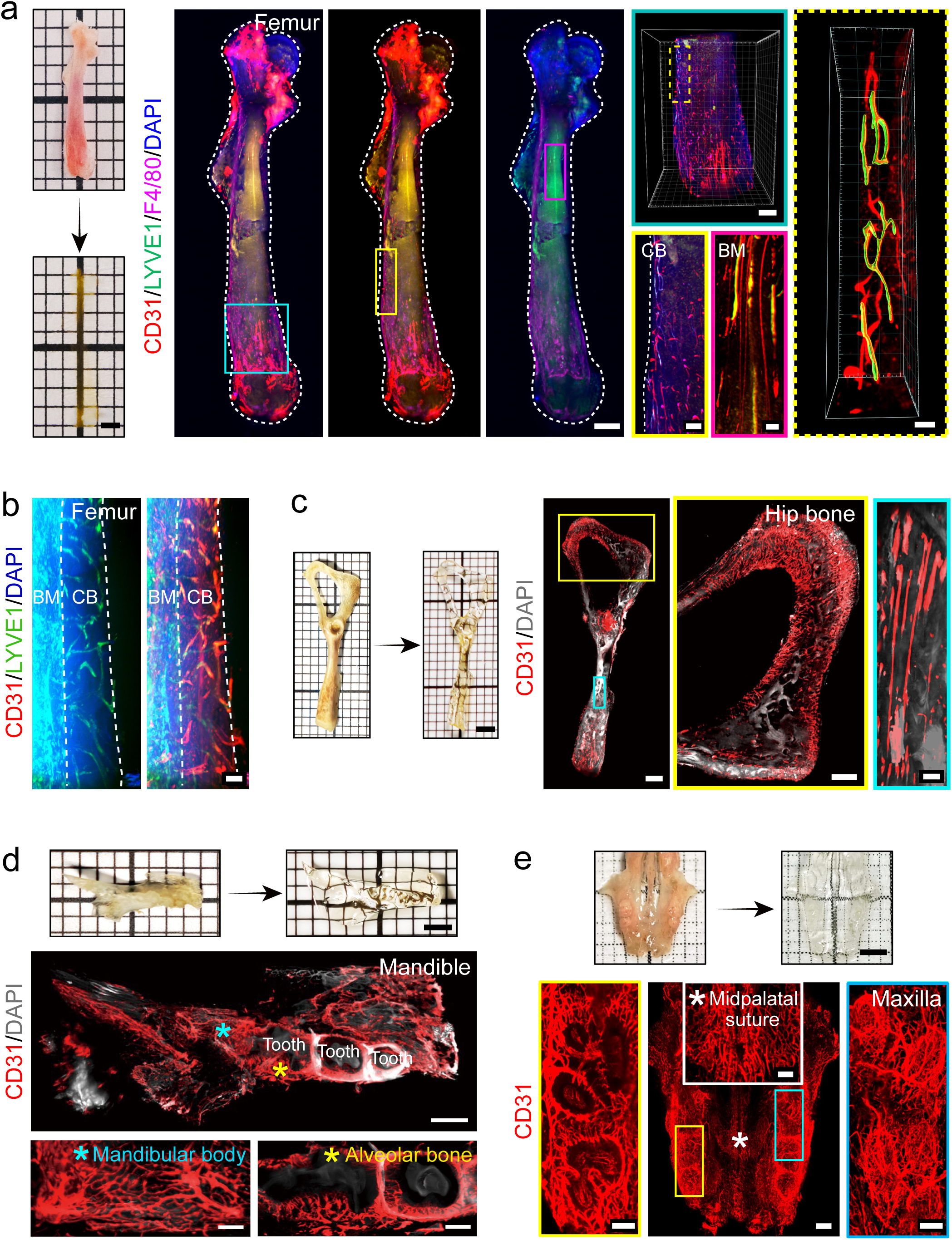
Light-sheet imaging for lymphatic vessels and blood vessels in various bones after tissue clearing. **a**, Images of intact mouse tibia prior to and post tissue-clearing. Panoptic light-sheet images showing vascular network immunolabeled by CD31 (red), lymphatic vessels immunolabeled by LYVE1 (green), and macrophages immunostained by F4/80 (violet), and nuclei labeled by DAPI (blue). Insets displaying the distribution patterns of signals in cortical bone and bone marrow with a surface rendering. Scale bars: left photo, 2 mm; middle panoptic images, 1000 µm; upper inset, 500 µm; lower left inset, 200 µm; lower right inset, 100 µm; right inset, 100 µm. BM, bone marrow; CB, cortical bone. **b**, Representative images of the cortical bone of a murine femur labeled with CD31 (red), LYVE1 (green), and DAPI (blue). Scale bar: 50 µm. BM, bone marrow; CB, cortical bone. **c**, Mouse hip bone prior to and post tissue-clearing. Panoptic light-sheet image of a hip bone stained by CD31 (red) and DAPI (grey). Insets showing high-magnification images of selected areas of interest in the hip bone. Scale bars: left photo, 2 mm; middle panoptic image, 800 µm; middle inset, 500 µm; right inset, 50 µm. **d**, Mouse mandible prior to and post tissue-clearing. Panoptic light-sheet image of a mandible stained by CD31 (red) and DAPI (grey). Insets showing high-magnification images of the mandibular body and alveolar bone. Scale bars: upper photo, 2 mm; middle panoptic image, 500 µm; lower insets, 200 µm. **e**, Mouse maxilla prior to and post tissue-clearing. Representative panoptic image of an intact maxilla stained with CD31 (red). Insets showing the high-resolution images of the area of interest. Scale bars: upper photo, 2 mm; lower left inset, 50 µm; lower middle panoptic image, 500 µm; lower middle inset, 50 µm; lower right inset, 50 µm.

Lymphatic vessels are delicate and sparsely distributed structures that are particularly difficult to visualize in bone using conventional imaging approaches. The protocol established in this study therefore provides a powerful tool for lymphatic vessel visualization and investigation. Consistent with our previous findings^31^, LYVE1-positive lymphatic vessels were observed in multiple intact murine bones **(Fig. 2a,b**, **Fig. 3a,b)**. As a subset of macrophages also expresses LYVE1, resulting in signal interference within the bone marrow, femoral samples were further processed with additional F4/80 immunolabeling to mitigate macrophage-derived LYVE1 signals **(Fig. 3a)**. This approach enabled high-resolution visualization of the spatial colocalization of CD31 and LYVE1 signals within the bone marrow, allowing precise delineation of lymphatic vessel localization **(Fig. 3a)**. Moreover, samples from transgenic mice with endogenic fluorescence can also be processed using this protocol to perform light-sheet imaging, as confirmed by the 3D images of intact bones from the lymphatic vessel reporter mouse line LYVE1-EGFP^31^. In addition to long bones, we also analyzed vascular network architectures in other irregular skeletal elements, including the hip bone, mandible, and maxilla **(Fig. 3c-e)**. Notably, after clearing, high-resolution visualization of vascular networks was achieved within both the jawbones and teeth **(Fig. 3d,e)**, highlighting the utility of this approach for studies of craniofacial skeletal biology.

Using Imaris File Converter, raw microscopy datasets were converted into .ims files and subsequently processed in Imaris for three-dimensional visualization **(Fig. 4a-d)**. This pipeline allows surface or filament rendering of the endothelial system, spot-based analysis of perivascular, other associated cell populations **(Fig. 4e)**, and systematic quantitative analysis **(Fig. 4f)**. Taking the mouse mandible as an example, we provide a detailed illustration of the complete image analysis workflow. Following processing with this pipeline and multicolor immunostaining, intact light-sheet microscopy datasets were obtained for both young and aged mandibles **(Extended Data Fig. 3)**. Regions of interest were then subjected to high-resolution scanning, followed by surface and spot rendering analyses, enabling clear visualization of vascular networks and macrophages at single-cell resolution **(Fig. 5a)**. Quantitative analyses based on these datasets revealed a significant age-associated reduction in blood and lymphatic vessels, accompanied by a marked expansion of macrophages in aged alveolar bones **(Fig. 5b)**. Detailed, step-by-step analyses of whole-tissue or organ-scale datasets are provided in **Supplementary Videos 6-8**, facilitating reproducibility of this workflow and enabling readers to obtain the anticipated results using this protocol.

**Fig. 4.**
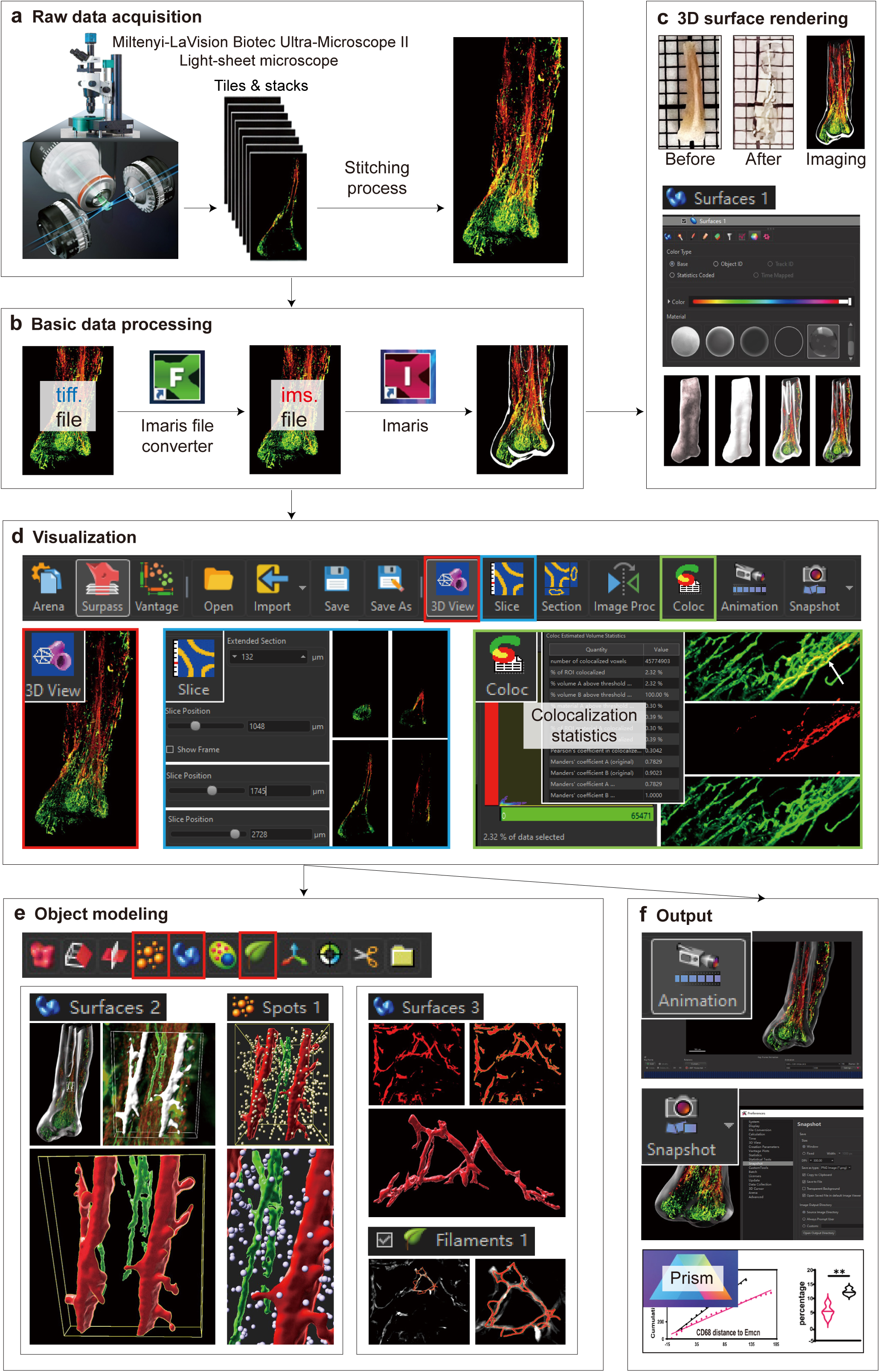
Pipeline for data processing and image analysis using Imaris software. **a**, Acquisition of raw image data using light-sheet microscopy with tiles and stacks scanning modes. Image stitching was performed using the bundled software. **b**, Basic data processing workflow starts with the file conversion from tiff. files to ims. files using the Imaris file converter for subsequent 3D modeling. **c.** 3D surface rendering of the cleared bone tissue and various rendering modalities available in Imaris. **d**, Visualization techniques including global 3D views, orthographic slice views, and colocalization analysis of multi-channel fluorescence signals. **e**, Object modeling utilizing specific modules, including Surfaces for volumetric structures, Spots for point-like objects, and Filaments for tracing tubular networks. **f**, Data output and quantification. Data are exported as animations (video) or high-resolution snapshots with customizable dimensions and pixels. Quantitative analysis is subsequently performed based on the modeled objects.

**Fig. 5.**
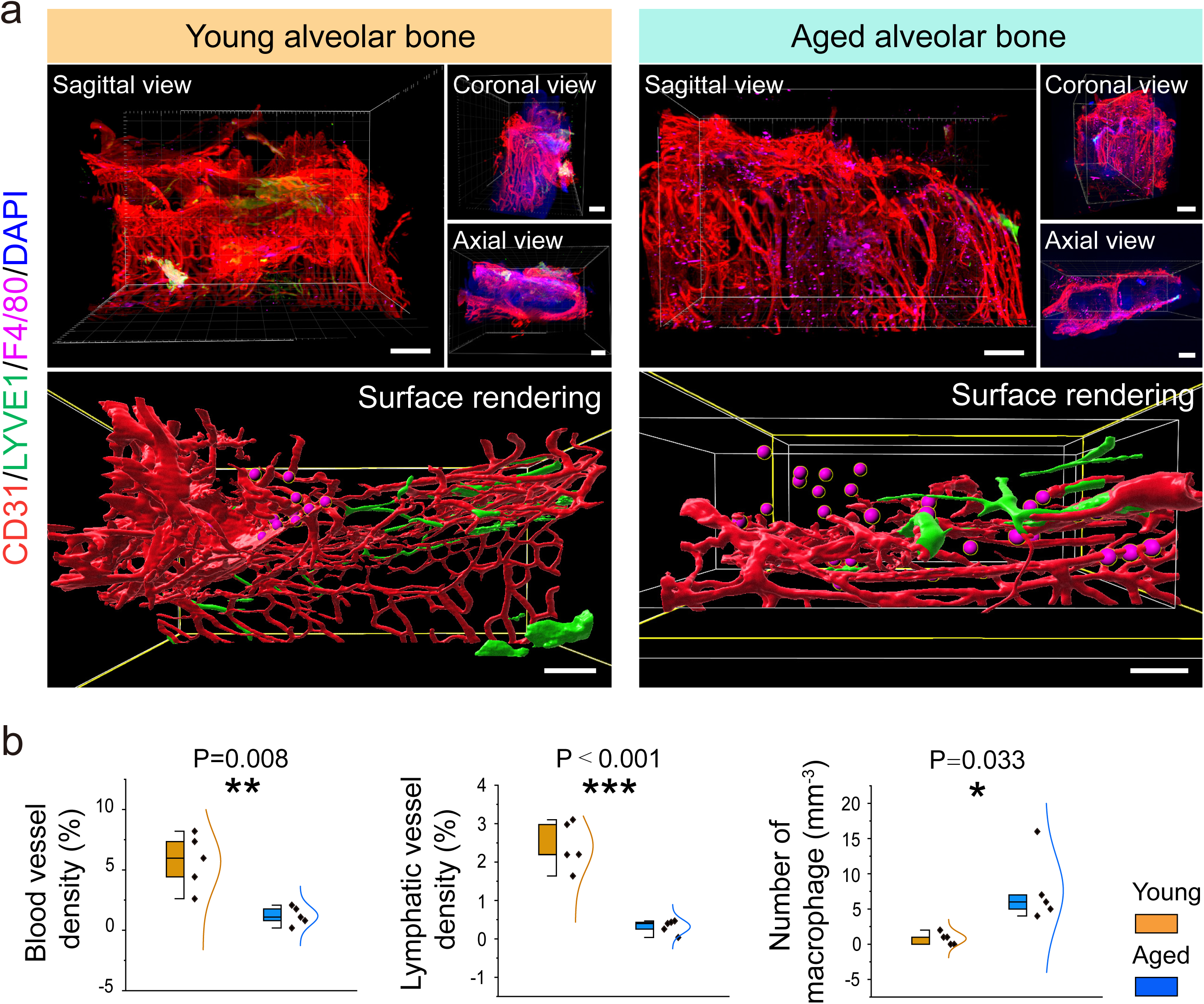
Light-sheet imaging and data analysis for the vascular system and macrophages in cleared alveolar bones. **a**, 3D images of young and aged mice alveolar bones stained with CD31 (red), LYVE1 (green), F4/80 (violet), and DAPI (grey) in different directions. Surface and spot rendering of the vascular system and macrophages in young and aged mice alvelolar bones. Scale bars: cropping, 300 µm; rendering, 150 µm. **b**, Quantitative analyses of vessels, lymphatic vessels, and macrophages in young and aged mice alveolar bones. Data represents mean ± s.d., p-value derived from two-tailed unpaired t tests (*p < 0.05, **p < 0.01, and ***p < 0.001).

## Data availability

The raw data underlying all graphical figures presented in this study are provided in the Data Source Files.

## Author contributions

Z.D., Y.S., J.C. and A.P.K. performed experiments. Z.D., Y.S. and H.L. analyzed the data. Z.D. and J.C. contributed to the preparation and writing of the manuscript. C.L. and J.C. commented on the manuscript. A.P.K. conceived and developed the method, designed experiments, provided funding support, and reviewed the manuscript.

## Acknowledgments

A.P.K is supported by Medical Research Council (CDA: MR/P02209X/1), European Research Council (StG: metaNiche, 805201), and Ministry of Education (MOE) Singapore: Academic Research Funds (#024983-00001, #025277-00026). J.C. is supported by National Natural Science Foundation of China (Nos. 82422021, 82270961) and Sichuan Provincial Health Commission (24QNMP015).

## Legends to Figures and Tables

**Extended Data Fig. 1 Light-sheet 3D and slice view images showing microenvironments in murine bones.**

**a**, Representative cropping images of tibial bone marrow labeled with CD31 (red), LYVE1 (green), F4/80 (violet), and DAPI (blue). Slice views demonstrating antibody penetration throughout multiple layers in the cropping volume consisting of 115 layers with a total thickness of 193 µm. Scale bars: 200 µm. BM, bone marrow.

**b**, Light-sheet images of a whole murine maxilla showing multicolor signals, including α-SMA (red) and CD31 (green) with high-resolution insets of molar regions. Slice view images showing antibody penetration throughout multiple layers in this maxilla consisting of 435 layers with a total thickness of 1053 µm. Scale bars: maxilla, 500 µm; molars, 50 µm; slices, 500 µm; right insets, 50 µm.

**Extended Data Fig. 2 Evaluation of samples digested by collagenase A and operation process of tibia dissection.**

**a**, Representative 3D images of mouse tibia from digestion and non-digestion groups labeled with CD31 (yellow) and DAPI (blue) with high-resolution insets of bone marrow regions. H&E staining of tibial sections from two groups. Scale bars: overview, 1000 µm; insets and H&E staining images,100 µm.

**b**, Representative photos showing the mouse tibia before and after removal of surrounding soft tissues.

**Extended Data Fig. 3 Light-sheet imaging of young and aged mice mandibles.**

Representative panoptic images of young and aged mice mandibles stained with CD31 (red), LYVE1 (green), F4/80 (violet), and DAPI (grey). Insets showing the high-resolution images of areas of interest. Scale bars: panoptic images, 1000 µm; alveolar bone, 200 µm; mandibular body, 100 µm.

**Table 1:** Troubleshooting advice

**Extended Data Table 1:** Primary and secondary antibodies details

**Extended Data Table 2:** Workstation configuration

**Supplementary video 1:** A whole murine maxilla, immune-stained with Endomucin, shows blood vessels in the intact bone tissue.

**Supplementary video 2:** A whole murine mandible, immune-stained with Endomucin, shows blood vessels in the intact bone tissue.

**Supplementary video 3:** A whole murine mandible, immune-stained with Endomucin and CD31, shows the type H vessels in the intact bone tissue.

**Supplementary video 4:** Panoptic multicolor immunolabeling for blood vessels and arteries on an intact murine tooth.

**Supplementary video 5:** An intact murine tibia labeled for blood vessels (Endomucin), cytotoxic T cells (CD8α), and nuclei (DAPI) with surface rendering.

**Supplementary video 6:** Animation showcasing the sequential progression of the image analysis.

**Supplementary video 7:** In-depth coverage of basic 3D analysis techniques with Imaris software.

**Supplementary video 8:** Detailed exploration of surface rendering processes.

## References

1 Belle, M. et al. Tridimensional Visualization and Analysis of Early Human Development. Cell 169, 161–173.e112 (2017).

2 Ding, Y. et al. Multiscale light-sheet for rapid imaging of cardiopulmonary system. JCI Insight 3, e121396 (2018).

3 Salwig, I. & Spitznagel, B. Imaging lung regeneration by light sheet microscopy. Cell Biol. 155, 271–277 (2021).

4 Di Giovanna, A. P., Tibo, A. & Silvestri, L. Whole-Brain Vasculature Reconstruction at the Single Capillary Level. Sci. Rep. 8, 12573 (2018).

5 Zhao, S. et al. Cellular and Molecular Probing of Intact Human Organs. Cell 180, 796–812.e19 (2020).

6 Neckel, P. H., Mattheus, U., Hirt, B., Just, L. & Mack, A. F. Large-scale tissue clearing (PACT): Technical evaluation and new perspectives in immunofluorescence, histology, and ultrastructure. Sci. Rep. 6, 34331 (2016).

7 Prahst, C., Ashrafzadeh, P. & Mead, T. Mouse retinal cell behaviour in space and time using light sheet fluorescence microscopy. Elife 9, e49779 (2020).

8 Todorov, M. I. et al. Machine learning analysis of whole mouse brain vasculature. Nat. Methods 17, 442–449 (2020).

9 Ramasamy, S. K. et al. Blood flow controls bone vascular function and osteogenesis. Nat. Commun. 7, 13601 (2016).

10 Zhu, X. et al. Ultrafast optical clearing method for three-dimensional imaging with cellular resolution. Proc. NatI. Acad. Sci. U. S. A. 116, 11480–11489 (2019).

11 Zhu, J. & Yu, T. MACS: Rapid Aqueous Clearing System for 3D Mapping of Intact Organs. Adv. Sci. (Weinh) 7, 1903185 (2020).

12 Davis, A. S. et al. Characterizing and Diminishing Autofluorescence in Formalin-fixed Paraffin-embedded Human Respiratory Tissue. J. Histochem. Cytochem. 62, 405–423 (2014).

13 Piña, R. et al. Ten Approaches That Improve Immunostaining: A Review of the Latest Advances for the Optimization of Immunofluorescence. Int. J. Mol. Sci. 23, 1426 (2022).

14 Kusumbe, A. P., Ramasamy, S. K., Starsichova, A. & Adams, R. H. Sample preparation for high-resolution 3D confocal imaging of mouse skeletal tissue. Nat. Protoc. 10, 1904–1914 (2015).

15 Chen, J., Lippo, L., Labella, R., Tan, S. L. & Marsden, B. D. Decreased blood vessel density and endothelial cell subset dynamics during ageing of the endocrine system. EMBO J. 40, e105242 (2021).

16 Chen, J. & Sivan, U. High-resolution 3D imaging uncovers organ-specific vascular control of tissue aging. Sci. Adv. 7, eabd7819 (2021).

17 Ramasamy, S. K., Kusumbe, A. P., Wang, L. & Adams, R. H. Endothelial Notch activity promotes angiogenesis and osteogenesis in bone. Nature 507, 376–380 (2014).

18 Kusumbe, A. P. et al. Age-dependent modulation of vascular niches for haematopoietic stem cells. Nature 532, 380–384 (2016).

19 Kusumbe, A. P., Ramasamy, S. K. & Adams, R. H. Coupling of angiogenesis and osteogenesis by a specific vessel subtype in bone. Nature 507, 323–328 (2014).

20 Ni, Y. et al. Deep imaging of LepR^+^ stromal cells in optically cleared murine bone hemisections. Bone Res. 13, 6 (2025).

21 Chieco, P., Jonker, A., De Boer, B. A., Ruijter, J. M. & Van Noorden, C. J. Image cytometry: protocols for 2D and 3D quantification in microscopic images. Prog. Histochem. Cytochem. 47, 211–333 (2013).

22 Chung, K. et al. Structural and molecular interrogation of intact biological systems. Nature 497, 332–337 (2013).

23 Griffini, P., Smorenburg, S. M., Verbeek, F. J. & van Noorden, C. J. Three-dimensional reconstruction of colon carcinoma metastases in liver. J. Microsc. 187, 12–21 (1997).

24 Snippert, H. J., Schepers, A. G., Delconte, G., Siersema, P. D. & Clevers, H. Slide preparation for single-cell-resolution imaging of fluorescent proteins in their three-dimensional near-native environment. Nat. Protoc. 6, 1221–1228 (2011).

25 Greenbaum, A. & Chan, K. Y. Bone CLARITY: Clearing, imaging, and computational analysis of osteoprogenitors within intact bone marrow. Sci. Transl. Med. 9, eaah6518 (2017).

26 Jing, D. et al. Tissue clearing of both hard and soft tissue organs with the PEGASOS method. Cell Res. 28, 803–818 (2018).

27 Chakraborty, T. & Driscoll, M. K. Light-sheet microscopy of cleared tissues with isotropic, subcellular resolution. Nat. Methods 16, 1109–1113 (2019).

28 Singh, A. et al. Angiocrine signals regulate quiescence and therapy resistance in bone metastasis. JCI insight 4, 125679 (2019).

29 Neu, C. P., Novak, T., Gilliland, K. F., Marshall, P. & Calve, S. Optical clearing in collagen- and proteoglycan-rich osteochondral tissues. Osteoarthritis Cartilage 23, 405–413 (2015).

30 Sdobnov, A. Y. et al. Recent progress in tissue optical clearing for spectroscopic application. Spectrochim. Acta. A Mol. Biomol. Spectrosc. 197, 216–229 (2018).

31 Biswas, L. et al. Lymphatic vessels in bone support regeneration after injury. Cell 186, 382–397.e24 (2023).

32 Shi, Y. et al. Vascular and Lymphatic Dysregulation via Non-EndoMT Col2a1 Signaling in Bisphosphonate-Related Osteonecrosis of the Jaw. bioRxiv, 09.27.678999 (2025).

33 Liu, H. et al. Lymphatic-immune interactions in the musculoskeletal system. Front. immunol. 16, 1578847 (2025).

34 Kumar, N., Saraber, P., Ding, Z. & Kusumbe, A. P. Diversity of Vascular Niches in Bones and Joints During Homeostasis, Ageing, and Diseases. Front. immunol. 12, 798211 (2021).

35 Yu, T., Zhu, J., Li, D. & Zhu, D. Physical and chemical mechanisms of tissue optical clearing. iScience 24, 102178 (2021).

36 Power, R. M. & Huisken, J. A guide to light-sheet fluorescence microscopy for multiscale imaging. Nat. Methods 14, 360–373 (2017).

37 Sato, Y. & Miyawaki, T. Quick visualization of neurons in brain tissues using an optical clearing technique. Anat. Sci. Int. 94, 199–208 (2019).

38 Messal, H. A. & Almagro, J. Antigen retrieval and clearing for whole-organ immunofluorescence by FLASH. Nat. Protoc. 16, 239–262 (2021).

39 Ertürk, A. et al. Three-dimensional imaging of solvent-cleared organs using 3DISCO. Nat. Protoc. 7, 1983–1995 (2012).

40 Dodt, H. U. et al. Ultramicroscopy: three-dimensional visualization of neuronal networks in the whole mouse brain. Nat. Methods 4, 331–336 (2007).

41 Kolesová, H., Čapek, M., Radochová, B., Janáček, J. & Sedmera, D. Comparison of different tissue clearing methods and 3D imaging techniques for visualization of GFP-expressing mouse embryos and embryonic hearts. Histochem. Cell Biol. 146, 141–152 (2016).

42 Wang, Q., Liu, K., Yang, L., Wang, H. & Yang, J. BoneClear: whole-tissue immunolabeling of the intact mouse bones for 3D imaging of neural anatomy and pathology. Cell Res. 29, 870–872 (2019).

43 Ueda, H. R. & Ertürk, A. Tissue clearing and its applications in neuroscience. Nat. Rev. Neurosci. 21, 61–79 (2020).

44 Susaki, E. A. et al. Advanced CUBIC protocols for whole-brain and whole-body clearing and imaging. Nat. Protoc. 10, 1709–1727 (2015).

45 Yang, B. et al. Single-cell phenotyping within transparent intact tissue through whole-body clearing. Cell 158, 945–958 (2014).

46 Kubota, S. I. et al. Whole-Body Profiling of Cancer Metastasis with Single-Cell Resolution. Cell Rep. 20, 236–250 (2017).

47 Utagawa, K. et al. Three-dimensional visualization of neural networks inside bone by Osteo-DISCO protocol and alteration of bone remodeling by surgical nerve ablation. Sci. Rep. 13, 4674 (2023).

48 Yi, Y. et al. Mapping of individual sensory nerve axons from digits to spinal cord with the transparent embedding solvent system. Cell Res. 34, 124–139 (2024).

49 Gorelashvili, M. G., Heinze, K. G. & Stegner, D. Optical Clearing of Murine Bones to Study Megakaryocytes in Intact Bone Marrow Using Light-Sheet Fluorescence Microscopy. Methods Mol. Biol. 1812, 233–253 (2018).

50 Rindone, A. N. & Liu, X. Quantitative 3D imaging of the cranial microvascular environment at single-cell resolution. Nat. Commun. 12, 6219 (2021).

51 Li, W., Germain, R. N. & Gerner, M. Y. High-dimensional cell-level analysis of tissues with Ce3D multiplex volume imaging. Nat. Protoc. 14, 1708–1733 (2019).

52 Mai, H. et al. Whole-body cellular mapping in mouse using standard IgG antibodies. Nat. Biotechnol. 42, 617–627 (2024).

53 Ou, Z. et al. Achieving optical transparency in live animals with absorbing molecules. Science. 385, eadm6869 (2024).

54 Mertens, T. F. & Liebheit, A. T. MarShie: a clearing protocol for 3D analysis of single cells throughout the bone marrow at subcellular resolution. Nat. Commun. 15, 1764 (2024).

