## Supplementary material for "Rapid immunostaining and high-resolution three-dimensional light-sheet microscopy of intact calcified tissues": Table 1

**Table 1: Troubleshooting advice**

| **Step** | **Problem** | **Possible reason** | **Solution** |
| --- | --- | --- | --- |
| 1 | **Excessive muscle tissue imaging and uneven internal bone tissue imaging** | 1. The muscle tissues are not thoroughly cleaned | Thoroughly remove the muscle from the surface of the bone tissue and carry out appropriate collagenase digestion |
|  |  | 2. The periosteum is not thoroughly cleaned | Carefully scrape off the periosteum from the surface of the bone tissue |
| 4 | **Suboptimal decalcification effect or uneven decalcification** | 1. The samples have insufficient contact with the decalcifying solution | Fill the decalcifying solution to near the capacity of the tube and ensure that all regions of bone tissue are adequately immersed in the liquid during rotation |
|  |  |  | Do not use too small a tube during decalcification, as it may lead to incomplete decalcification. |
|  |  | 2. EDTA is not fully dissolved, leading to insufficient solution concentration | Thoroughly stir for an adequate duration when preparing the EDTA solution |
|  |  | 3. Incorrect pH | Adjust the pH of the decalcifying solution to 7.4 |
| 6, 7 | **Suboptimal bleaching effect** | 1. The samples are not thoroughly dehydrated before the bleaching step | Dehydrate the sample strictly following the requirements of solution concentration gradients and time intervals |
|  |  | 2. Insufficient H_2_O_2_ solution concentration | The commercially available H_2_O_2_ solutions typically have concentrations of 20-30%, and they need to be diluted to 5%. Please make sure to perform the necessary and correct calculations |
|  |  |  | Store H_2_O_2_ away from light and at 4 °C, as it degrades over time, resulting in a reduction in solution concentration |
| 11 | **Suboptimal collagenase digestion effect** | 1. The collagenase becomes inactive due to prolonged storage | Ensure that a fresh collagenase solution is prepared each time |
|  |  | 2. Enzymatic digestion conducted at excessively high or low incubation temperatures | Carefully adjust the water bath temperature to 37 °C |
| 1, 5, 6, 7, 8, 9, 10, 16, 18, 20, 21 and 23 | **Weak fluorescence staining/no signals** | 1. The periosteum and muscle are not completely removed | Thoroughly remove the muscle from the surface of the bone tissue and carry out appropriate collagenase digestion |
|  |  | 2. EDTA is not thoroughly washed off, interfering with immunostaining | After decalcification, make sure to thoroughly wash away EDTA based on the step requirements |
|  |  | 3. Prolonged bleaching can damage surface antigens of the tissue | Strictly control the bleaching time within 2 hours |
|  |  | 4. Improper permeabilization leads to inadequate antibody penetration | Strictly adhere to the formulation of the permeabilization solution and incubation time |
|  |  | 5. Low antibody concentration | Adjust the antibody concentration based on the recommended dilution ratio |
|  |  | 6. Compromised antibody quality | Pay attention to the antibody's expiration date, use new antibodies if required |
|  |  |  | Avoid freeze-thaw cycles, spin down the antibody and mix it gently before use |
|  |  | 7. Suboptimal antibody incubation | Strictly adhere to the incubation time and the incubation temperature of 37 °C |
|  |  | 8. Fluorescence quenching | Avoid excessive dehydration and conduct dehydration at 4 °C to minimize the influences of ethanol and methanol on samples |
|  |  | 9. Photobleaching | Ensure protection from light exposure during and after the immunostaining process |
|  |  | 10. Poor sample transparency | Please refer to the solutions below |
| 16, 17, 18 and 19 | **Noisy and nonspecific fluorescent particles** | 1. The antibodies are not thoroughly washed during the staining process | Carefully follow the required washing steps for both primary and secondary antibodies |
|  |  | 2. Precipitates and aggregates in the secondary antibody | Avoid repeated freeze-thaw cycles, and spin down the secondary antibody before use |
|  |  | 3. The samples become dry during the staining incubation process | Maintain sample moisture throughout all steps |
| 6, 7, 14, 16 and 18 | **High background staining** | 1. Suboptimal bleaching, remaining pigmentation and hematoma in tissue | Please refer to the solutions above |
|  |  | 2. Suboptimal sample blocking | Strictly adhere to the required blocking time and the incubation temperature of 37 °C |
|  |  | 3. Very high concentration of the antibody used | Reduce the antibody concentration |
| 2, 6, 7, 20, 21 and 23 | **Poor sample transparency** | 1. Too much hematin within the bone marrow | Conduct perfusion fixation with 4% PFA before bone dissection, and increase the bleaching period |
|  |  | 2. Inadequate dehydration after immunostaining | Strictly follow the requirements of solution concentration gradients and time intervals |
|  |  | 3. Inadequate clearing incubation for large bone samples | Prolong the incubation time of the sample in the clearing medium |
| 23, 28 and 29 | **Bubbles disrupt the imaging quality during image acquisition** | 1. Bubbles appearing within bone tissue, including structures like the alveolar nerve canal | Carefully fill the empty cavities with the clearing medium by needle without damaging the tissues |
|  |  | 2. Bubbles appearing during sample attachment to the adaptor holder | Carefully use a 23 G needle with a 2 mL syringe to remove the sample and attach again |
|  |  | 3. Bubbles appearing on the sample’s surface when placed into the imaging solution | Slowly place the sample into the imaging solution to prevent the formation of bubbles |
