## Supplementary figures and images for "Rapid immunostaining and high-resolution three-dimensional light-sheet microscopy of intact calcified tissues"

### Figure S1

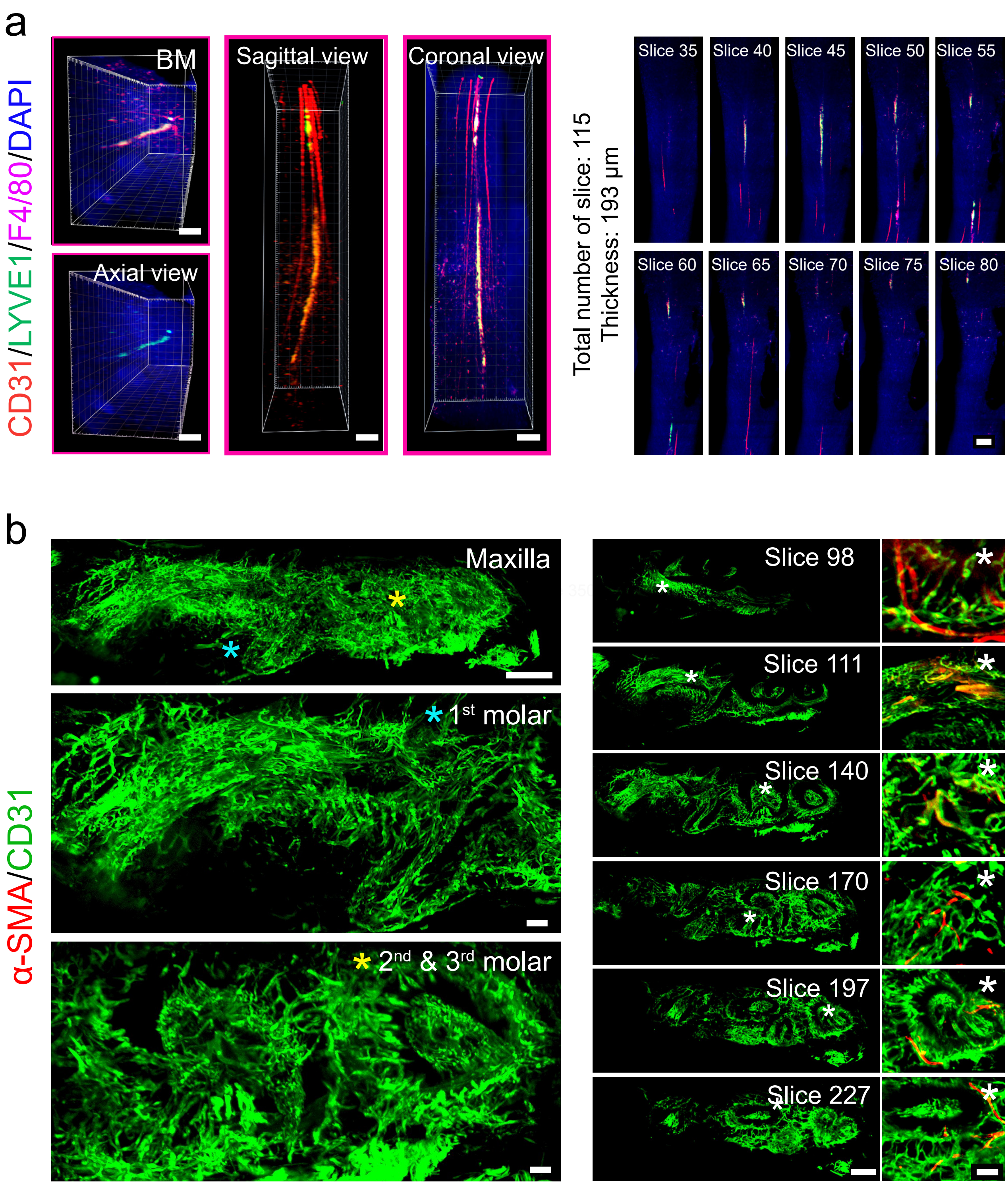


Extended Data Figure 1

### Figure S2

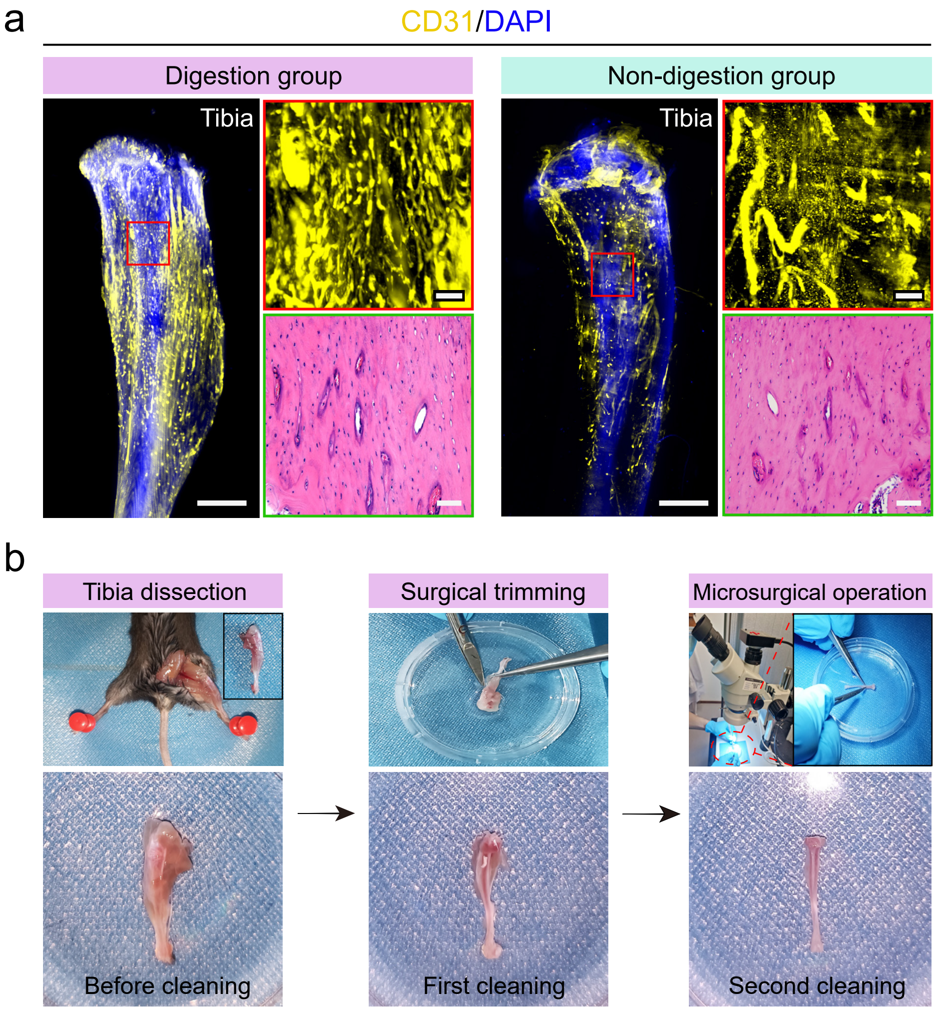


#### Extended Data Figure 2

### Figure S3

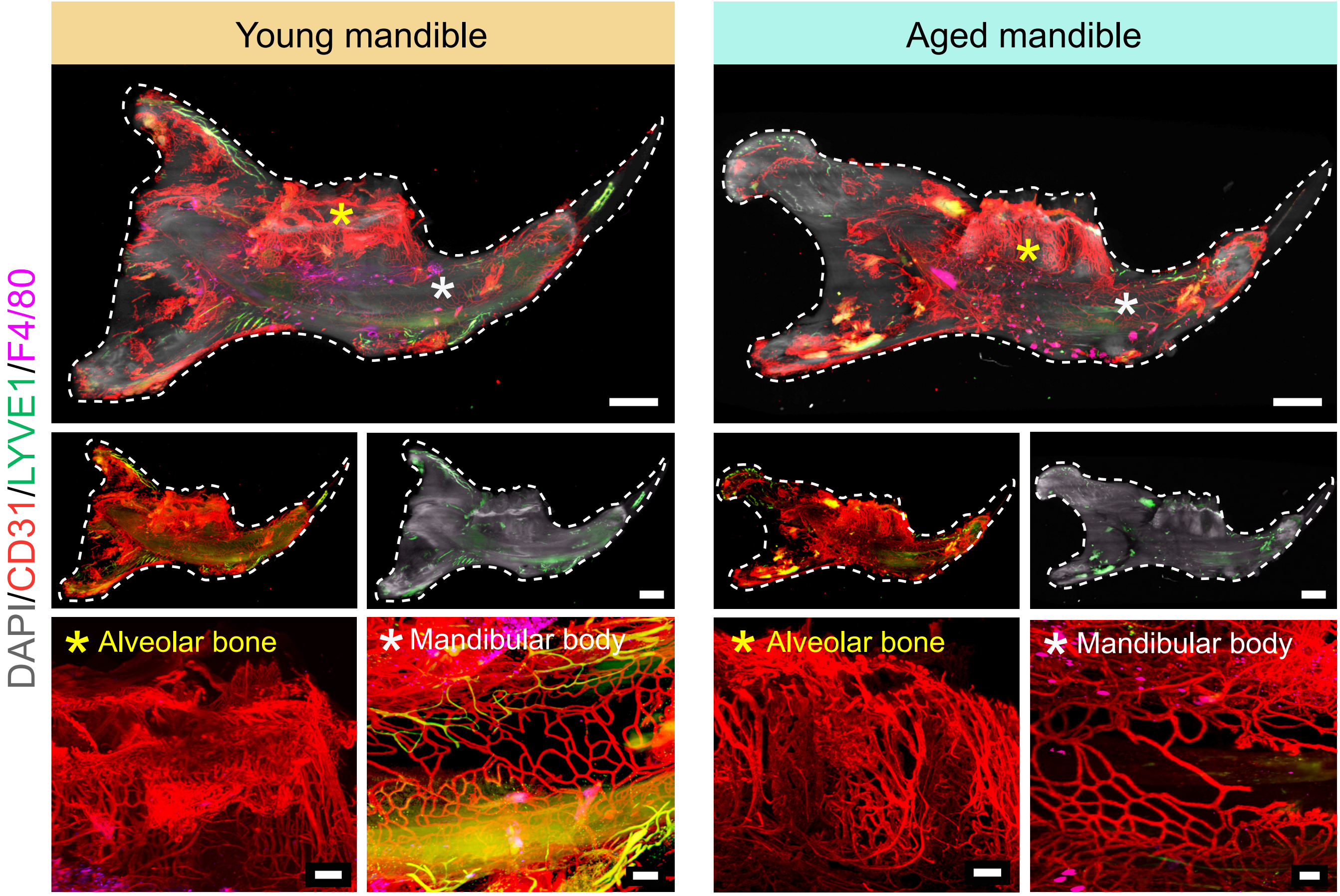


Extended Data Figure 3
